# Real-time biomedical knowledge synthesis of the exponentially growing world wide web using unsupervised neural networks

**DOI:** 10.1101/2020.04.03.020602

**Authors:** Tyler Wagner, Samir Awasthi, Gayle Wittenberg, AJ Venkatakrishnan, Dan Tarjan, Anuli Anyanwu-Ofili, Andrew Badley, John Halamka, Christopher Flores, Najat Khan, Rakesh Barve, Venky Soundararajan

**Affiliations:** nference, inc., One Main St, East Arcade, Suite 400, Cambridge, MA 02142, USA; Janssen pharmaceutical companies of Johnson & Johnson (J&J), USA; Mayo Clinic, Rochester, MN 55905, USA

## Abstract

Decoding disease mechanisms for addressing unmet clinical need demands the rapid assimilation of the exponentially growing biomedical knowledge. These are either inherently unstructured and non-conducive to current computing paradigms or siloed into structured databases requiring specialized bioinformatics. Despite the recent renaissance in unsupervised neural networks for deciphering unstructured natural languages and the availability of numerous bioinformatics resources, a holistic platform for real-time *synthesis* of the scientific literature and seamless triangulation with deep omic insights and real-world evidence has not been advanced. Here, we introduce the n*f*erX platform that makes the highly unstructured biomedical knowledge computable and supports the seamless visual *triangulation* with statistical inference from diverse structured databases. The n*f*erX platform will accelerate and amplify the research potential of subject-matter experts as well as non-experts across the life science ecosystem (https://academia.nferx.com/).

---

The n*f*erence cloud-based software platform (n*f*erX) enables dynamic inference from 45 quadrillion possible conceptual associations that synthesize over 100 million documents scraped from the published world wide web, and this is continuously updated as new material is published online. The platform supports visual triangulation of insights via statistical enrichments from nearly 50,000 curated collections of structured databases, with the diseases, biomolecules, drugs, and cells & tissues collections loaded by default. A hypergeometric test is used to capture the overlap between the knowledge synthesis results and each of these enrichment sets. The sources include all freely accessible literature integrated into a Core Corpus as well as distinct sub-corpora that provide contextual lenses into PubMed, preprints, clinical trials, SEC filings, patents, grants, media, company websites, etc. The collections include curated ontologies or statistical inference applied to molecular data (e.g. genomics, bulk and single cell RNA-seq, proteomics) and real-world data (e.g. FDA adverse event reports, clinical trial outcomes, epidemiology, clinical case reports). Here, we describe how the n*f*erX platform^1^ can enable data science driven decision-making via a pair of illustrative applications -- (i) biopharmaceutical lifecycle management across conventionally siloed therapeutic areas, and (ii) connecting clinical pathophysiology to molecular profiling for a rapidly evolving pandemic.

As described previously, the n*f*erX platform adeptly identifies well-known and emerging associations embedded in the biomedical literature using two key metrics^2^: *local context score* and *global context score* (**Figure 1A**). The local context score is based on two significant improvements over the traditional pointwise mutual information (PMI)^3^. First, we extend the PMI-based strength of association to biomedical concepts that can be constructed by a logical combination of proximal phrases, e.g. “EGFR-positive” AND “non-small cell lung cancer”. This effectively makes the number of biomedical concepts that can be queried unbounded. Moreover, we extend the traditional PMI notion, which is unable to capture the word-distance between co-occurring terms, using “exponential masking” to meaningfully account for the distance between co-occurring terms, captured by “score decay” in n*f*erX. Our experimental studies show several measures for which our local score method outscores traditional PMI metrics *(unpublished results)*. To compute the global context score we use an unsupervised neural network with dependency parsing to generate over 300 million biomedical multi-word phrase vectors, and leverage word2vec^4^ to compute the cosine distance between these phrase vectors is projected in a 300-dimensional space. Previous studies of word embeddings provide a heuristic to extend their unigram technique to specify multi-word terms apriori^4^, and cui2vec relies on a curated set of phrases with associated vectors^5^. However, our examination indicates that relying on such occurrence frequency or curated data is not as exhaustive as our approach for generating a complete list of biomedical terms of interest, in particular when there are many low frequency or emerging concepts *(unpublished results)*. We have also devised a novel method to compute vectors “on demand” for very low frequency phrases, for which pre-computing vectors is exorbitantly expensive.

**Figure 1.**
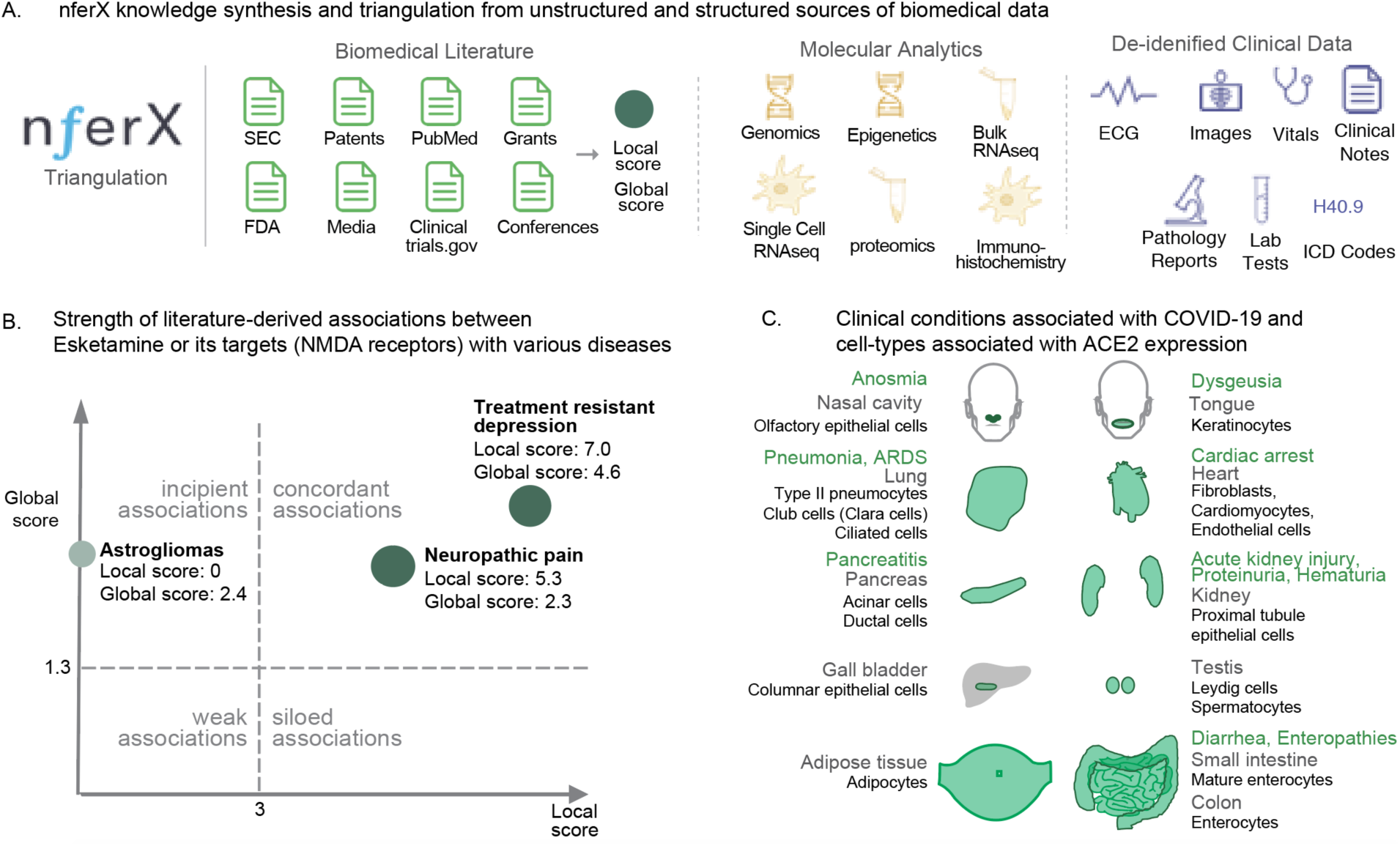
Illustrative applications of the nferX platform: Biopharma lifecycle management and clinical-molecular linkage. **(A)** Schematic diagram highlighting computation of literature-derived association scores. **(B)** A lifecycle management study of esketamine recapitulated well-known associations to neuropathic pain and treatment resistant depression, as well as an emerging association between the drug’s target NMDA receptors and astrogliomas. The predicted astrogliomas association to NMDA receptors was prospectively validated by a subsequent study^6^. **(C)** Linking the emerging clinical pathophysiology of COVID-19 patients with organs expressing the ACE2 viral receptor.

As an application of how researchers can use the combination of local and global context to add credence to existing hypotheses and identify novel associations, n*f*erX was used to investigate potential indications for the label expansion of esketamine, an NMDA receptor antagonist recently approved for treatment-resistant depression (TRD). A hypothesis-free analysis was performed using the platform to quantify associations between esketamine, its targets (NR2A, NR2B), and all possible indications {<u>n</u>*<u>f</u>*<u>erX link</u>}. n*f*erX automatically populated the synonyms for the key biomedical entities such as genes, as an “expanded query”. The platform correctly recapitulated relationships between esketamine and well-known indications, such as treatment resistant depression (local score = 7) and neuropathic pain (local score = 5.3), but also identified novel associations such as neuro-oncogenesis (e.g. global score of 2.4 between ‘astrogliomas’ and ‘NMDA receptors’; <u>n</u>*<u>f</u>*<u>erX link</u>) (**Figure 1B**). This link to neuro-oncogenesis was subsequently confirmed^6^, approximately 3 months after our analysis was initially performed. Such a prospective validation adds credence to further interrogate esketamine for other emerging indications identified as having significant global score to NMDA receptors (e.g. neurogenic inflammation, global score = 2.4, local score = 0.9, <u>n</u>*<u>f</u>*<u>erX link</u>).

As another application, the n*f*erX platform enabled the rapid, comprehensive literature-based and multi-omic profiling of ACE2^7^, the putative receptor of SARS-CoV2 (**Figure 1C**). We found that tongue keratinocytes and olfactory epithelial cells of the nasal cavity are potentially novel ACE2-expressing cell populations. Clinically this aligns with the known routes of transmission through droplet spread and viral attachment within oral/nasal mucosa^8,9^. Also this may explain the altered sense of smell and taste in otherwise asymptomatic COVID-19+ individuals^8,9^. Next the very high rates of viral pneumonitis in infected patients with ground glass infiltrates on chest imaging^10^ are a logical sequelae of infection given the expression of ACE2 in type-2 pneumocytes, club cells and ciliated cells of the lung^7^. The copious ACE2 expression in various gastrointestinal (GI) cell types emphasizes the recent reports of diarrhea^11^ and signs/symptoms of enteropathy that are seen clinically, and may also explain the occurrence of fecal shedding that persists post-recovery^12^. Applying n*f*erX to identify incipient associations to ACE2 in the disease collection highlights diabetic renal disease, cardiorenal syndrome, and nephropathy hypertension (each with a global score of 3.3 to ACE2), as well as the enrichment sets of renal insufficiency, heart failure and kidney diseases {<u>n</u>*<u>f</u>*<u>erX link</u>}. These emerging insights from n*f*erX are consistent with diabetes mellitus and chronic kidney disease being identified as the leading mortality indicators for the elderly COVID-19+ patients^13^. This may also provide a pathophysiological rationale as to why some COVID patients experience complications like acute kidney injury^14^, proteinuria, hematuria, or myocarditis with associated rise in troponins^15^. Along these lines, it will be of interest to see if cases of SARS-CoV2 induced orchitis occurs in COVID-19 patients, given ACE2 expression in cells of the testes^7^, and the high n*f*erX local score of 4.8 for the orchitis-infertility association {<u>n</u>*<u>f</u>*<u>erX link</u>}.

The n*f*erX data science platform will help researchers generate insights via holistic triangulation of structured and unstructured data at an unprecedented scale. The full clinical potential of the unsupervised neural networks that power this platform will be realized when they are applied towards automated de-identification and synthesis of the unstructured physician notes that dominate the Electronic Health Records (EHRs). To enable such seamless real-world insight triangulation with the wealth of published biomedical knowledge, a privacy-preserving federated architecture that exports aggregate statistical inferences while retaining the primary de-identified data within the academic medical center’s span of control is needed. Such a platform can truly propel clinical research and biopharmaceutical development into the digital era.

## References

1. The nferX platform. https://academia.nferx.com/

2. Park, J. et al. Recapitulation and Retrospective Prediction of Biomedical Associations Using Temporally-enabled Word Embeddings. doi:10.1101/627513 (2019).

3. Holzinger, A., Yildirim, P., Geier, M. & Simonic, K.-M. Quality-Based Knowledge Discovery from Medical Text on the Web. Intelligent Systems Reference Library 145–158 (2013) doi:10.1007/978-3-642-37688-7_7.

4. Tomas Mikolov, Ilya Sutskever, Kai Chen, Greg Corrado, Jeffrey Dean. Distributed Representations of Words and Phrases and their Compositionality. https://arxiv.org/abs/1310.4546.

5. Andrew L. Beam, Benjamin Kompa, Allen Schmaltz, Inbar Fried, Griffin Weber, Nathan P. Palmer, Xu Shi, Tianxi Cai, Isaac S. Kohane. Clinical Concept Embeddings Learned from Massive Sources of Multimodal Medical Data. https://arxiv.org/abs/1804.01486.

6. Barria, A. Dangerous liaisons as tumour cells form synapses with neurons. Nature vol. 573 499–501 (2019).

7. Venkatakrishnan, A. J. et al. Knowledge synthesis from 100 million biomedical documents augments the deep expression profiling of coronavirus receptors. doi:10.1101/2020.03.24.005702 (2020).

8. COVID-19. https://www.entuk.org/categories/covid-19.

9. AAO-HNS: Anosmia, Hyposmia, and Dysgeusia Symptoms of Coronavirus Disease. American Academy of Otolaryngology-Head and Neck Surgery https://www.entnet.org/content/aao-hns-anosmia-hyposmia-and-dysgeusia-symptoms-coronavirus-disease (2020).

10. Kanne, J. P., Little, B. P., Chung, J. H., Elicker, B. M. & Ketai, L. H. Essentials for Radiologists on COVID-19: An Update—Radiology Scientific Expert Panel. Radiology 200527 (2020) doi:10.1148/radiol.2020200527.

11. Gu, J., Han, B. & Wang, J. COVID-19: Gastrointestinal manifestations and potential fecal-oral transmission. Gastroenterology (2020) doi:10.1053/j.gastro.2020.02.054.

12. Xu, Y. et al. Characteristics of pediatric SARS-CoV-2 infection and potential evidence for persistent fecal viral shedding. Nature Medicine (2020) doi:10.1038/s41591-020-0817-4.

13. Bhatraju, P. K. et al. Covid-19 in Critically Ill Patients in the Seattle Region — Case Series. New England Journal of Medicine (2020) doi:10.1056/nejmoa2004500.

14. Cheng, Y. et al. Kidney disease is associated with in-hospital death of patients with COVID-19. Kidney International (2020) doi:10.1016/j.kint.2020.03.005.

15. Inciardi, R. M. et al. Cardiac Involvement in a Patient With Coronavirus Disease 2019 (COVID-19). JAMA Cardiology (2020) doi:10.1001/jamacardio.2020.1096.

